# An Enzyme-Linked Immunosorbent Assay (ELISA)-based Activity Assay for AMP-Activated Protein Kinase (AMPK)

**DOI:** 10.1101/2024.08.20.608877

**Authors:** Trezze P Nguyen, Yang Liu

## Abstract

AMP-activated protein kinase (AMPK) is the master regulator of cellular and organismal energy homeostasis, playing an essential role in modulating metabolism as well as other cellular processes. Efforts have been made to discover pharmacological modulators of AMPK activity because of their therapeutic potentials against various diseases. Measuring AMPK activity in vitro is a fundamental step for testing AMPK activators and inhibitors. Here we report an enzyme-linked immunosorbent assay (ELISA)-based AMPK activity assay with simple steps and high sensitivity, which represents an alternative in-house method to the traditional radioactive method as well as other techniques that require special reagents and commercial reagent kits.

## Introduction

AMP-activated protein kinase (AMPK) plays an essential role in maintaining cellular and organismal energy homeostasis [1]. It is activated when the cellular ATP levels decrease and AMP and ADP levels increase [1]. Upon activation, AMPK helps restore cellular energy balance through phosphorylating various downstream substrates which leads to the inhibition of many energy-consuming anabolic pathways, such as protein synthesis, fatty acid synthesis, etc., and the activation of energy-producing catabolic pathways, such as glucose uptake and glycolysis, fatty acid oxidation, etc. [2]. Besides its function in controlling metabolic pathways, AMPK also regulates autophagy process, mitochondrial homeostasis, lysosomal homeostasis, DNA repair, etc. [1, 3]. Given the key role of AMPK in controlling many cellular processes especially metabolism, the development of pharmacological modulators of AMPK (especially activators) as therapeutic strategies for treating various diseases, particularly the metabolic diseases and syndromes, has attracted great interest [4].

AMPK is a heterotrimeric complex containing one catalytic subunit (α) and two regulatory subunits (β and γ). In mammalian cells, there are two α isoforms (α1 and α2), two β isoforms (β1 and β2) and three γ isoforms (γ1, γ2 and γ3), which gives rise to 12 different heterotrimeric combinations. Certain heterotrimers may be dominantly expressed in specific tissues or cell types [1]. AMPK can be activated through phosphorylation of a threonine residue on the activation loop of the α subunit by upstream kinases such as LKB1 and CaMKK2 [1]. In addition, AMPK can also be activated allosterically. For example, AMP binds to the γ subunit which leads to allosteric activation of AMPK [5]. Moreover, many direct allosteric activators of AMPK discovered in recent years bind to the ADaM (Allosteric Drug and Metabolite) site which locates at the interface between α and β subunits [4]. Some of these activators have shown great therapeutic potential in treating metabolic diseases and syndromes such as diabetes and NASH (non-acholic nonalcoholic steatohepatitis) [6, 7].

Measuring AMPK kinase activity in vitro is a critical step for identifying small molecule AMPK activators and inhibitors as well as investigating regulatory mechanism of AMPK activation. The classic methods involve utilizing ^32^P- or ^33^P-ATP in the kinase reaction and measuring the incorporation of ^32^P or ^33^P into the kinase substrate peptide. In recent years, different non-radioactive methods have been proposed including several FRET (Fluorescence Resonance Energy Transfer)-based assays [8, 9] that measures the amount of phosphorylated substrate in a kinase reaction as well as other assays that measure the usage of ATP in a kinase reaction [10]. In these assays, special reagents such as fluorescent dye or biotin-labeled substrate peptides or/and antibodies and commercial reagent kits are usually needed. Here we report an ELISA-based AMPK activity assay with simple steps and high sensitivity using relatively common reagents. In this study, we apply this method to calculate kinase kinetics of recombinant AMPK complex as well as test the effects of known AMPK small molecule modulators. This method represents a convenient, in-house assay for AMPK kinase activity and a useful tool for the researchers in the AMPK research field.

### Materials and Methods

### Reagents

Recombinant AMPK α1β1γ1 was expressed in *E*.*coli* before purification and activation by CamKK2 as described previously [9]. AMPK substrate, SAMS peptide (sequence: HMRSAMSGLHLVKRR), and AMP solution were purchased from SignalChem Biotech (Richmond, BC, Canada; cat# S07-58 and cat# A46-09-500, respectively). Phospho-SAMS peptide (sequence:HMRSAMpSGLHLVKRR) was synthesized by Alan Scientific (Gaithersburg, MD, USA). ATP solution was purchased form New England Biolabs (Ipswich, MA, USA; cat# P0756). Anti-phospho-Acetyl CoA Carboxylase (Ser79) antibody and Horseradish peroxidase (HRP)-conjugated donkey anti-rabbit IgG antibody were purchased from Millipore Sigma (Burlington, MA, USA; cat# 07-303 and cat# AP182P, respectively). One-Step Ultra TMB ELISA substrate solution was purchased from Thermo Fisher Scientific (Waltham, MA, USA; cat# 34028). AMPK inhibitor, compound C (cat# AOB2281), and AMPK activators, compound 991 (cat# AOB8150) and PF-739 (cat# AOB33584), were purchased from AOBIOUS (Gloucester, MA, USA).

### AMPK activity assay using ELISA

AMPK kinase reactions are performed in PCR tubes. For each kinase reaction (20 μl), 4 μl of 5× kinase buffer (200mM Tris-HCl, pH7.4, 100mM MgCl_2_, 0.5mg/ml BSA and 0.25mM DTT), 50 μM AMP, 20 ng recombinant AMPK α1β1γ1, 0.2 μg/ul (112 μM) SAMS peptide, 500 μM ATP, and H_2_O (molecular biology level; to make the final volume 20 ul) are added. In some experiment, AMP is excluded in the reactions, and variable concentrations of AMPK, SAMS peptide and ATP are used as indicated in the text or figure legends. In the experiment that AMPK inhibitor (compound C) and AMPK activators (compound 991 and PF739) are tested, the stock solution of the inhibitor or activators (all dissolved in DMSO) with proper concentrations and DMSO (as control) were first diluted 200 times with H_2_O before 1 μl of these diluted solutions were added into each kinase reaction mixture to make the final concentration of DMSO 0.05% (v/v). ATP is added lastly to start the kinase reactions which will be carried out at room temperature for the period of time as indicated in the figure legends. If multiple reaction time points are studied as shown in Figure 1 (D), after reaching one desired reaction time point, 4.5 μl of the reaction mixture is taken out and mixed with 4.5 μl 50 mM EDTA to stop the kinase reaction, and this sample harvesting is repeated for the desired time points. If only one reaction period is used in the experiments, when reaching the time point, 20 ul 50 mM EDTA is added to the reaction mixture to stop the kinase reaction. To make the blank control without kinase reaction, kinase mixture without ATP is directly added to 20 ul 50 mM EDTA before ATP is added in at last. For the experiment that requires a standard curve for measuring the amount of phosphorylated SAMS peptide in each kinase reaction, a serial of peptide mixtures with a constant total peptide concentration of 200 ng/μl but contain different proportions of SAMS peptide and phospho-SAMS peptide were made in 1x kinase buffer. Specifically, a series of mixtures containing 0 ng/μl (0 pmole/20 μl), 2 ng/μl (21.34 pmole/20 μl), 4 ng/μl (42.69 pmole/20 μl),10 ng/μl (106.72 pmole/20 μl), 20 ng/μl (213.43 pmole/20 μl), 40 ng/μl, (426.86 pmole/20 μl) 80 ng/μl (853.72 pmole/20 μl), 120 ng/μl (1280.58 pmole/20 μl) of phospho-SAMS peptide (with variable amount of SAMS peptide to make the total concentration of peptide 200 ng/μl) were made and then 20 μl of these mixtures were mixed with 20 μl 50 mM EDTA. The stopped reaction mixtures including the blank and standard curve samples are further diluted 5 times with peptide coupling buffer (100 mM carbonate buffer, pH 9.6) before 5 ul of the diluted mixture from each reaction is added into one well of the Nunc Immobilizer Amino microplate (Thermo Fisher Scientific, Waltham, MA, USA; cat# 436013) filled with 95 ul of peptide coupling buffer. Microplate is putted either at 4°C overnight or at room temperature for 2 hours with gentle agitation to let the peptides from the reaction mixtures covalently coupled to the bottom surfaces of the plate wells. After the coupling process, the coupling mixtures are dumped out and the wells are washed three time with TBST (Tris-buffered saline with 0.1% (v/v) Tween 20; 300 μl/well/wash). The wells are then blocked with blocking buffer containing 1% (w/v) BSA in TBST (100 μl/well) for 1 hour at room temperature with gentle agitation. The blocking buffer is then dumped out and 70 μl of primary antibody solution containing anti-phospho-Acetyl-CoA Carboxylase (Ser79) antibody diluted in blocking buffer (1:200 dilution) is added into each well for 1 hour at room temperature. After the incubation with primary antibody, the plate is washed 6 times with TBST (300 μl/well/wash) before 100 μl of secondary antibody solution containing Horseradish peroxidase (HRP)-conjugated anti-rabbit IgG antibody diluted in blocking buffer (1:5000 dilution) is added into each well for another 1 hour at room temperature. The plate is then washed 6 times with TBST (300 μl/well/wash), and then 50 μl One-Step Ultra TMB ELISA substrate solution is added into each well for 10 to 20 min (plate is hidden from light during the incubation). When a desired blue color has appeared, 50 μl ELISA stop solution (0.16 M sulfuric acid) is added per well to stop the enzyme reaction. The plate is then read in a plate reader to measure optical density (OD) at 450 nm and 650 nm (as background wavelength correction). The value of OD 450 in each well is subtracted by the value of OD 650 before being used for further data analysis.

### Data analysis

The corrected OD450 value in each well from each kinase reaction is subtracted by the blank (or the values of the condition with 0 min reaction time or 0 μM substrate in some experiment) before plotted on the graph for comparation. For standard curve, only the values in the linear parts are used to establish the R^2^ value and the equation used to calculate the amount of phospho-SAMS from the OD value by linear regression analyzing function in GraphPad Prism 9.3.1. For Michaelis-Menten curves, Km are calculated using nonlinear regression - enzyme kinetics analyzing functions in GraphPad Prism 9.3.1. Graphs were made and statistics were analyzed with GraphPad Prism 9.3.1. Student’s t-test were used to compare differences between conditions.

## Results and Discussion

The aim of this work is to develop an ELISA-based assay to measure AMPK kinase activity through measuring the levels of phosphorylated SAMS peptide (an AMPK substrate peptide derived from Acetyl-CoA Carboxylase) after kinase reaction in vitro. In a kinase reaction performed in PCR tubes, AMPK phosphorylates SAMS peptide to generated phospho-SAMS peptide. And the amount of phospho-SAMS peptide generated within a certain period of reaction reflects the AMPK kinase activity. To quantify the relative levels of phospho-SAMS peptide in different reactions, the reactions mixtures are added into the wells of Nunc Immobilizer Amino microplate where the non-phosphorylated and phosphorylated SAMS peptides containing the nucleophilic primary amine at the N terminus and side chain of lysine are covalently conjugated to the bottom surface of microplate wells containing electrophilic groups. The levels of the phospho-SAMS peptide in each well are then detected by ELISA using anti-phospho-Acetyl-CoA Carboxylase (Ser79) antibody which recognizes the phospho-SAMS peptide. Within the range where the ELISA signal is not saturated, the relative higher OD value indicates more phospho-SAMS peptide generated due to higher AMPK kinase activity in the reaction. In the experimental process, the coupling of phospho-SAMS peptide to the surface of the microplate well is an essential step that can be affected by factors such as the kinase buffer of choice and may require optimization by the end users to determine how much of the kinase reaction mixture to be used for coupling. The experimental procedure listed in this study was optimized based on the kinase buffer we chose containing Tris-HCl and BSA, both of which contain primary amine and could be potentially coupled to the surface of the microplate wells, although we did not observe the interference from these two components on the experimental results we obtained.

Because it is crucial to establish the kinase reaction conditions to avoid signal saturation in the following ELISA step, we first tested different amount of AMPK in the kinase reaction (40 min reaction time) in the presence of AMP (50 μM) with saturated concentrations of substrate SAMS peptide (0.2 μg/μl or 112 μM) and ATP (500 μM) according to previous studies [11], and examined the OD value obtained from the latter ELISA step (**Fig.1A**). We found that ELISA signals remained in the linear range (R^2^= 0.9750) when up to 60 ng AMPK (∼ 21 nM) was used in the reaction. Based on this result, we chose the condition of 20 ng (∼6.9 nM) AMPK/reaction in the rest of this study. If the end users need to use a higher concentration of AMPK in the reaction, they may consider reducing the reaction time and test the ELISA signal saturation in their hands. To calculate the specific activity of AMPK based on the OD values, in the same experiment of Fig.1A, we also included a standard curve with various defined amounts of phospho-SAMS peptide/reaction (**Fig.1B**) and found that the signal remained in a linear range (R^2^= 0.9731) with up to 427 pmole phospho-SAMS peptide/reaction. Within this linear range, an equation, Y (OD value) = 0.002318*X (pmole of phospho-SAMS peptide/reaction) - 0.01645, which can be used to calculate the amount of phospho-SAMS peptide (pmole) in each reaction based on the OD value was established. Using this equation, we calculated the specific activity of AMPK in the reaction based on the results of the 20 ng AMPK/reaction (OD = 0.3724) in Fig.1A, and obtained 0.2097 pmole/ng/min which means that 0.2097 pmole SAMS peptide is phosphorylated by 1 ng of AMPK every min (**Fig.1C**). This specific activity value is in the same range as the value obtained in a previous study (∼0.7 pmole/ng/min) using radioactive method when recombinant AMPK α1β1γ1 complex which was purified and activated similarly as in this study was tested [12]. Given the fact that ELISA signals vary between different experiments, if specific activity needs to be calculated for one experiment, a standard curve need to be made in the same experiment to have accurate results. For the same reason, the OD values from different experiments should not be compared directly.

**Fig. 1.**
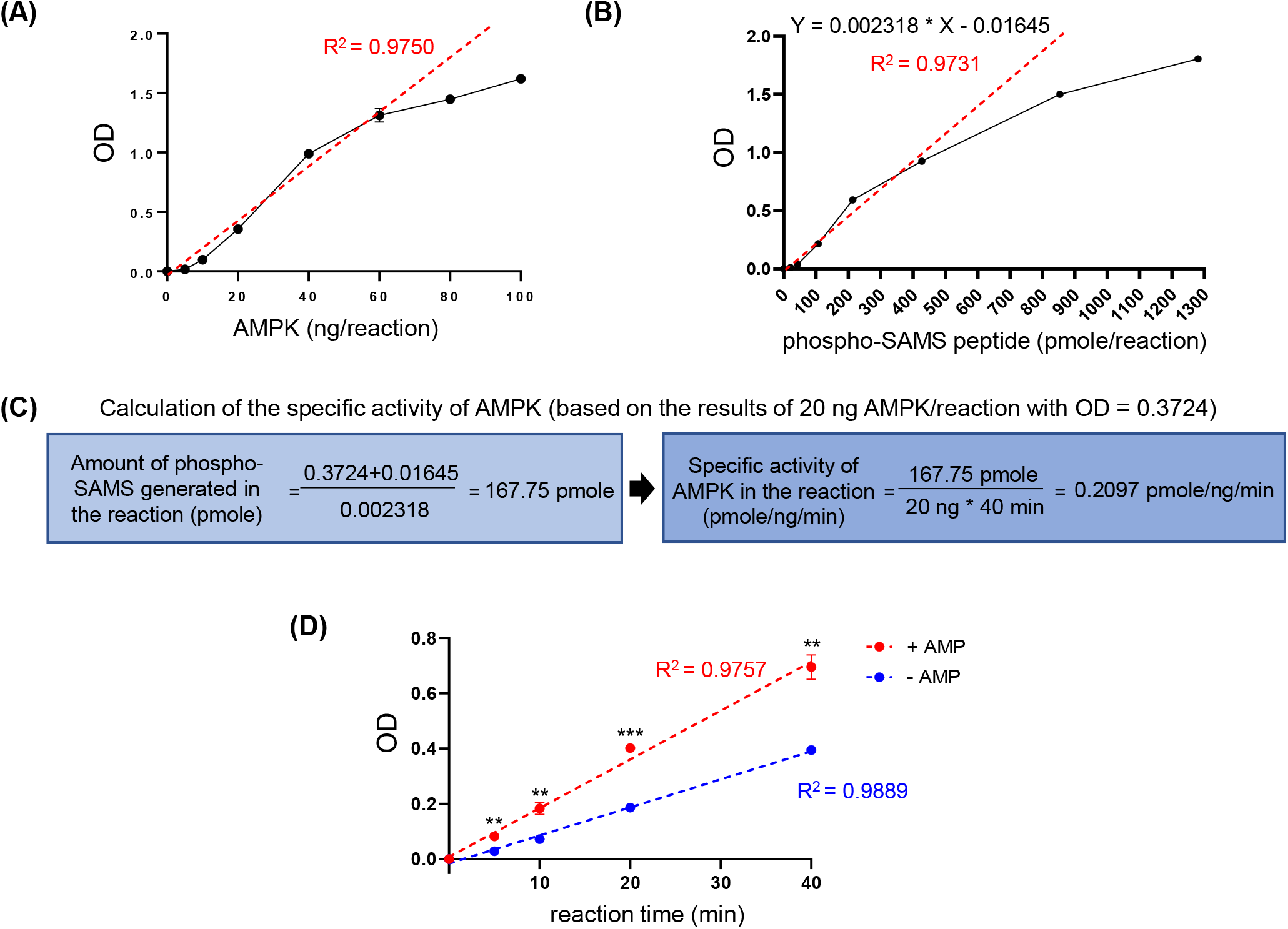
Validation of the proposed AMPK activity assay. **(A)** OD values from the ELISA following the kinase reactions (40 min reaction time) with different amount of AMPK (0, 5, 10, 20, 40, 60, 80 or 100 ng per reaction), n = 4. OD values from the condition of 0 ng AMPK was considered as blank and were subtracted from all the OD values in other conditions. R^2^ was calculated based on the data from conditions with 0, 5, 10, 20, 40 and 60 ng AMPK/reaction (red dotted line). **(B)** a standard curve established using different concentrations of phospho-SAMS peptide (0, 21.34, 42.69, 106.72, 213.43, 426.86, 853.72, or 1280.58 pmole/reaction), n = 3. OD values from the condition of 0 pmole/reaction was considered as blank and were subtracted from all the OD values in other conditions. R^2^ and the equation between Y (OD value) and X (pmole phospho-SAMS peptide/reaction) was established based on the data from the conditions with 0, 21.34, 42.69, 106.72, 213.43, 426.86 pmole phospho-SAMS peptide/reaction (red dotted line). (B) was done in the same experiment as shown in (A). **(C)** specific activity of AMPK based on the condition of 20 ng AMPK/reaction shown in (A) (OD=0.3724) was calculated using the equation established in (B). **(D)** OD values from the ELISA following 0 min, 5 min, 10 min, 20 min and 40 min of kinase reactions with 20 ng AMPK per reaction in the presence or absence of AMP, n = 3. OD values from the condition of 0 min reaction was considered as blank and were subtracted from all the OD values in other conditions. R^2^ was calculated based on the data from all the time points. ** *p* < 0.01, *** *p* < 0.001 when comparing the + AMP condition to the -AMP condition with the same reaction time by Student’s t-test. Data are mean ± s.e.m.

To further validate the method, we performed the kinase reaction (with 20 ng of AMPK) for different periods of time in the presence or absence of AMP before ELISA for determining the relative amounts of phospho-SAMS peptide generated in the reactions (**Fig.1D**). As expected, we observed a linear correlation between the reaction time and OD value (R^2^ = 0.9757 and 0.9889 for with AMP and without AMP conditions, respectively) with higher OD values observed in the reactions with the presence of AMP. These results above demonstrate the validity as well as the adequate accuracy and sensitivity of the method.

In addition, we also applied the method to determine the enzymatic kinetics of AMPK using different concentrations of substrates, ATP or SAMS peptide (**Fig.2A and B**). Michaelis-Menten curves were drawn based on the concentrations of the substrates in the reaction and the related OD values, and Km were calculated from the curves. From this experiment, the Km values we obtained were 25.27 μM for ATP and 15.68 μM for SAMS peptide, which are within a similar range as the values obtained in another study (26.04 μM for ATP and 26.67 μM for SAMS peptide) by radioactive method using recombinant AMPK α1β1γ1 complex purified and activated similarly as in this study [11]. These results further prove the reliability of this method.

**Fig. 2.**
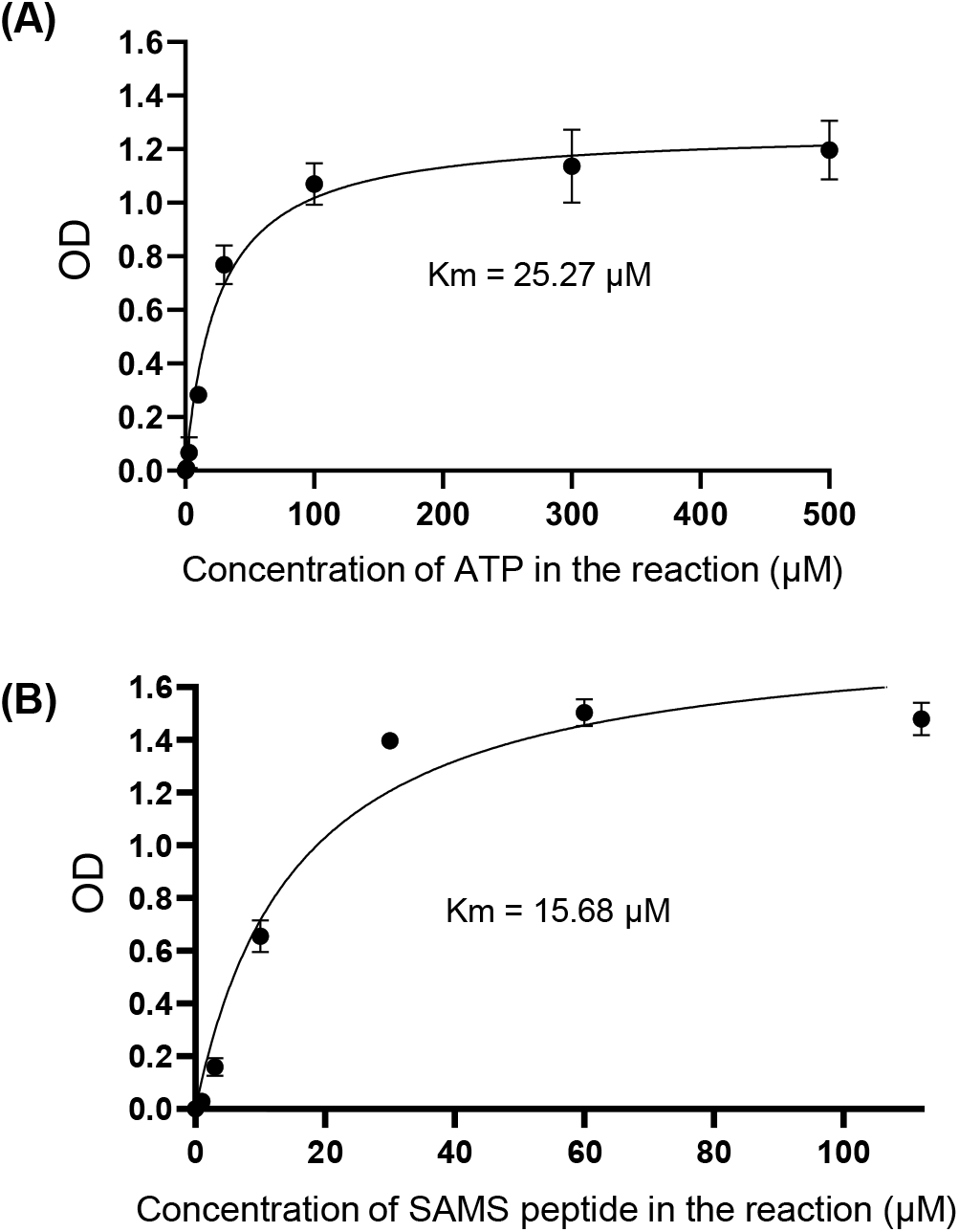
Enzymatic kinetics of AMPK determined using the proposed activity assay. **(A)** Michaelis-Menten curve based on the concentrations of the ATP (0, 1, 3, 10, 30, 100, 300 or 500 μM) in the reaction (40 min reaction time) and the related OD values, n = 3. Values from the condition of 0 μM ATP was considered as blank and were subtracted from all the OD values in other conditions. **(B)** Michaelis-Menten curve based on the concentrations of the SAMS peptide (0, 1, 3, 10, 30, 60, 112 μM) in the reaction (40 min reaction time) and the related OD values, n = 4. Values from the condition of 0 μM SAMS peptide was considered as blank and were subtracted from all the OD values in other conditions. Km values in (A) and (B) was calculated using nonlinear regression-enzyme kinetics analyzing functions in GraphPad Prism 9.3.1. Data are mean ± s.e.m.

Next, we employed the method to examine the effects of several known AMPK activity modulators including one inhibitor, compound C [13], and two direct allosteric activators, compound 991 [14, 15] and PF-739 [6], on AMPK kinase activity in the presence and absence of AMP in the reaction (**Fig.3**). As expected, compound C (5 μM) treatment significantly reduced AMPK kinase activity (by ∼ 64% and 69% in conditions with or without AMP, respectively), while compound 991 (250 nM) or PF-739 (250 nM) treatment significantly increased the AMPK kinase activity in both conditions with or without AMP (for compound 991, ∼ 3.0 and 1.8 fold increase in the conditions with or without AMP, respectively; for PF-739, ∼ 3.6 and 2.9 fold increase in the conditions with or without AMP, respectively). And in all the treatment conditions, the AMPK activity is higher with the presence of AMP in the reaction. The data demonstrated the feasibility of using this method to test or verify the effects of direct AMPK modulators in vitro.

**Fig. 3.**
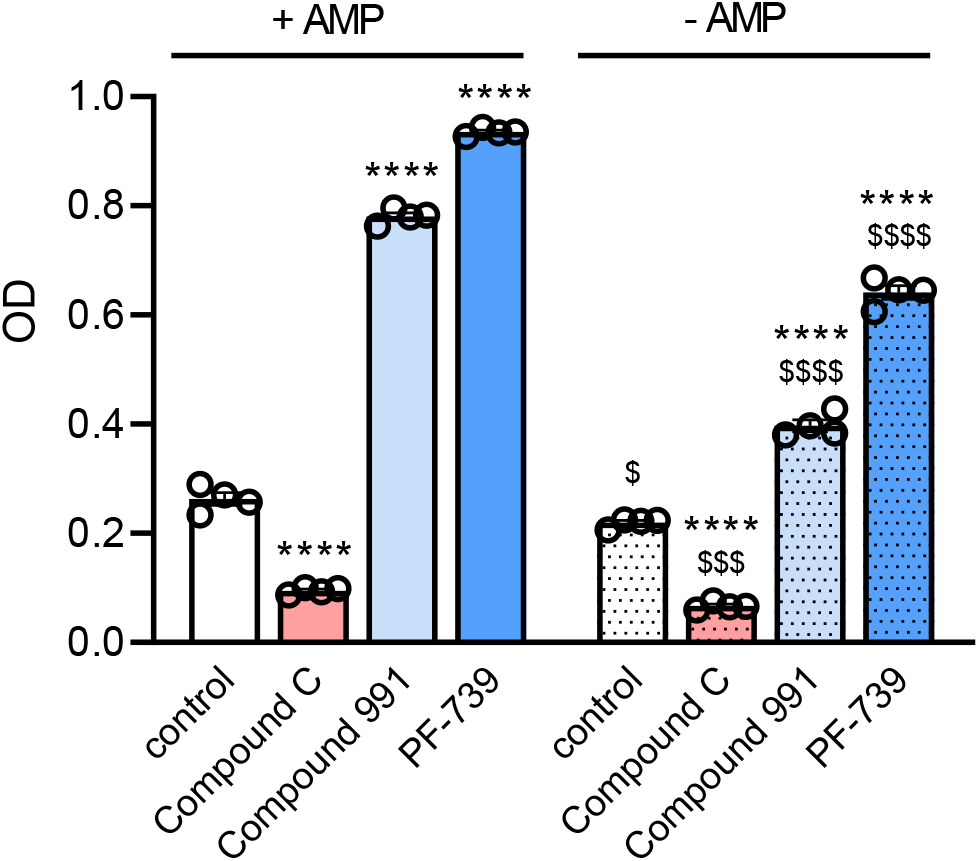
Effects of AMPK activity modulators tested using the proposed activity assay. Compound C (5 μM; an AMPK inhibitor), compound 991 (250 nM; an AMPK activator), PF-739 (250 nM; an AMPK activator) or DMSO control were added into the kinase reaction (40 min reaction time) in the presence or absence of AMP. OD values from the ELISA following the kinase reactions were measured and compared, n = 4. For each treatment condition, the OD value from 0 min reaction was determined and considered as blank, and was subtracted from the OD value of the experimental condition. **** *p* < 0.0001 when comparing to control condition with the same AMP availability in the reaction using Student’s t-test. $ *p* < 0.05, $$$ *p* < 0.001, $$$$ *p* < 0.0001 when comparing to the condition with the same modulator treatment but with the presence of AMP using Student’s t-test. Data are mean ± s.e.m.

## Conclusions

Our results denoted the great feasibility, reliability and sensitivity of the ELISA-based AMPK activity assay described in this study. This simple, in-house method represents an alternative to the traditional radioactive method as well as other techniques requiring special reagents and commercial reagent kits, and can be applied to identify and test AMPK modulators for pharmaceutical purposes.

## Acknowledgements

We thank Dr. Kei Sakamoto from University of Copenhagen for providing the recombinant AMPK complex as well as constructive discussion regarding the preparation of the manuscript.

## Abbreviations

AMP: adenosine monophosphate
AMPK: AMP-activated protein kinase
ADP: adenosine diphosphate
ADaM: Allosteric Drug and Metabolite
ATP: adenosine triphosphate
CamKK2: Calcium/calmodulin-dependent protein kinase kinase 2
ELISA: Enzyme-Linked Immunosorbent Assay
LKB1: Liver kinase B1
OD: optical density.

